# α-catenin links integrin adhesions to F-actin to regulate ECM mechanosensing and rigidity-dependence

**DOI:** 10.1101/2021.01.21.427565

**Authors:** Abhishek Mukherjee, Shay Melamed, Hana Damouny-Khoury, Malak Amer, Lea Feld, Elisabeth Nadjar-Boger, Michael P. Sheetz, Haguy Wolfenson

## Abstract

Both cell-cell and cell-matrix adhesions are regulated by mechanical signals, but the mechanobiological processes that mediate the crosstalk between these structures are poorly understood. Here we show that α- catenin, a mechanosensitive protein that is classically associated with cadherin-based adhesions, directly interacts with and regulates integrin adhesions. α-catenin is recruited to the edges of mesenchymal cells, where it interacts with F-actin. This is followed by mutual retrograde flow of α-catenin and F-actin from the cell edge, during which α-catenin interacts with vinculin within integrin adhesions. This interaction affects adhesion maturation, stress-fiber assembly, and force transmission to the matrix. In epithelial cells, α-catenin is present in cell-cell adhesions and absent from cell-matrix adhesions. However, when these cells undergo epithelial-to-mesenchymal transition, α-catenin transitions to cell-matrix adhesions, where it facilitates proper mechanosensing. This is highlighted by the ability of α-catenin-depleted cells to grow on soft matrices. These results suggest a dual role of α-catenin in mechanosensing, through both cell-cell and cell-matrix adhesions.

## Introduction

The ability of cells to sense and respond to their immediate extracellular environment affects the most fundamental cellular functions, such as survival, proliferation, and migration (Ringer et al., 2017; Iskratsch et al., 2014; Yap et al., 2018). This sensing ability relies on direct physical interactions with neighboring cells or with the extracellular matrix (ECM), which are facilitated by cell adhesion molecules – cadherins and integrins, respectively (Maiden and Hardin, 2011). The balance and transition between these two types of interactions can determine the state of a cell, such as in the process of epithelial to mesenchymal transition (EMT). However, the mechanisms of interplay between cell-cell and cell-matrix adhesions remain poorly understood.

Cell-cell contacts are classically mediated by adherens junctions (AJs), which are composed of transmembrane E-Cadherin molecules and the catenin family of proteins – p120-catenin, β-catenin, and α-catenin – that bind as a complex to the cytoplasmic tails of the cadherins (Pokutta and Weis, 2007). α-catenin is an actin-binding protein, and its ability to mediate the connection between cadherin and the actin cytoskeleton is vital for AJs, as actomyosin-based forces are required for stabilizing the cadherin-cadherin connection (Sarpal et al., 2019; Yonemura et al., 2010). Myosin II motors that operate on actin filaments create tension in α-catenin and cause structural changes in its C-terminal actin-binding domain (ABD), and in its M-domain (Thomas et al., 2013). Thus, α-catenin acts as a mechanosensory protein, since it is activated when forces are applied to it (Barry et al., 2014; Buckley et al., 2014). The M-domain recruits numerous adhesion-related proteins, including ZO-1, afadin1, α-actinin, and vinculin (Kobielak and Fuchs, 2004). Notably, α-actinin and vinculin are also important mediators of the connection between integrins and filamentous actin (F-actin) within focal adhesions (FAs) (Parsons et al., 2010; Wolfenson et al., 2013), which suggests a possible involvement of α-catenin in these structures. Indeed, previous studies have suggested a role for α-catenin outside of cell-cell adhesions (Sun et al., 2014; Vassilev et al., 2017; Wood et al., 2017; Piao et al., 2014). However, whether α-catenin plays a role in regulating cell-matrix adhesions and ECM mechanosensing is unknown.

Here we show that in mesenchymal cells, α-catenin is recruited to the cell edge, where it interacts with actin in regions devoid of α-actinin. It then undergoes retrograde flow together with F-actin towards the cell center, and interacts with vinculin within integrin adhesions. This interaction mediates adhesion maturation, enhances force transmission to the matrix, and drives the proper assembly of actin stress fibers. We find that while the loss of α-catenin is not sufficient to induce EMT on its own, it does play a role in rigidity-dependent EMT induced by transforming growth factor β (TGFβ). Importantly, post-EMT, α-catenin transitions to the edges of mesenchymal cells, where it facilitates mechanosensing. Moreover, the absence of the α-catenin–vinculin interaction causes mesenchymal cells to display impaired adhesion to the matrix. This results in aberrant mechanosensing of the matrix, and the transformation of cells to being rigidity-independent for growth.

## Results

### α-catenin localizes to the edges of fibroblast cells

Since α-catenin was shown to be recruited to mesenchymal cell edges (Wood et al., 2017), we postulated that it might play a role in regulating integrin adhesions that assemble in these regions. To test this, we first verified the recruitment of α-catenin to the cell edges of three different fibroblast cell lines: mouse embryonic fibroblasts (which henceforth will be referred to as MEFs), NIH3T3, and human foreskin fibroblasts (HFF). We plated the cells sparsely on fibronectin (FN)-coated coverslips to prevent the formation of cell-cell contacts, and fixed them after 15 minutes of spreading. We confirmed by immunostaining that in 50-60% of these cells α-catenin was localized in lamellipodial regions (typically presented as narrow stripes at the cell edge; ∼500 nm for MEF and NIH3T3, ∼1200 nm for HFF) (Fig. 1A).

**Figure 1.**
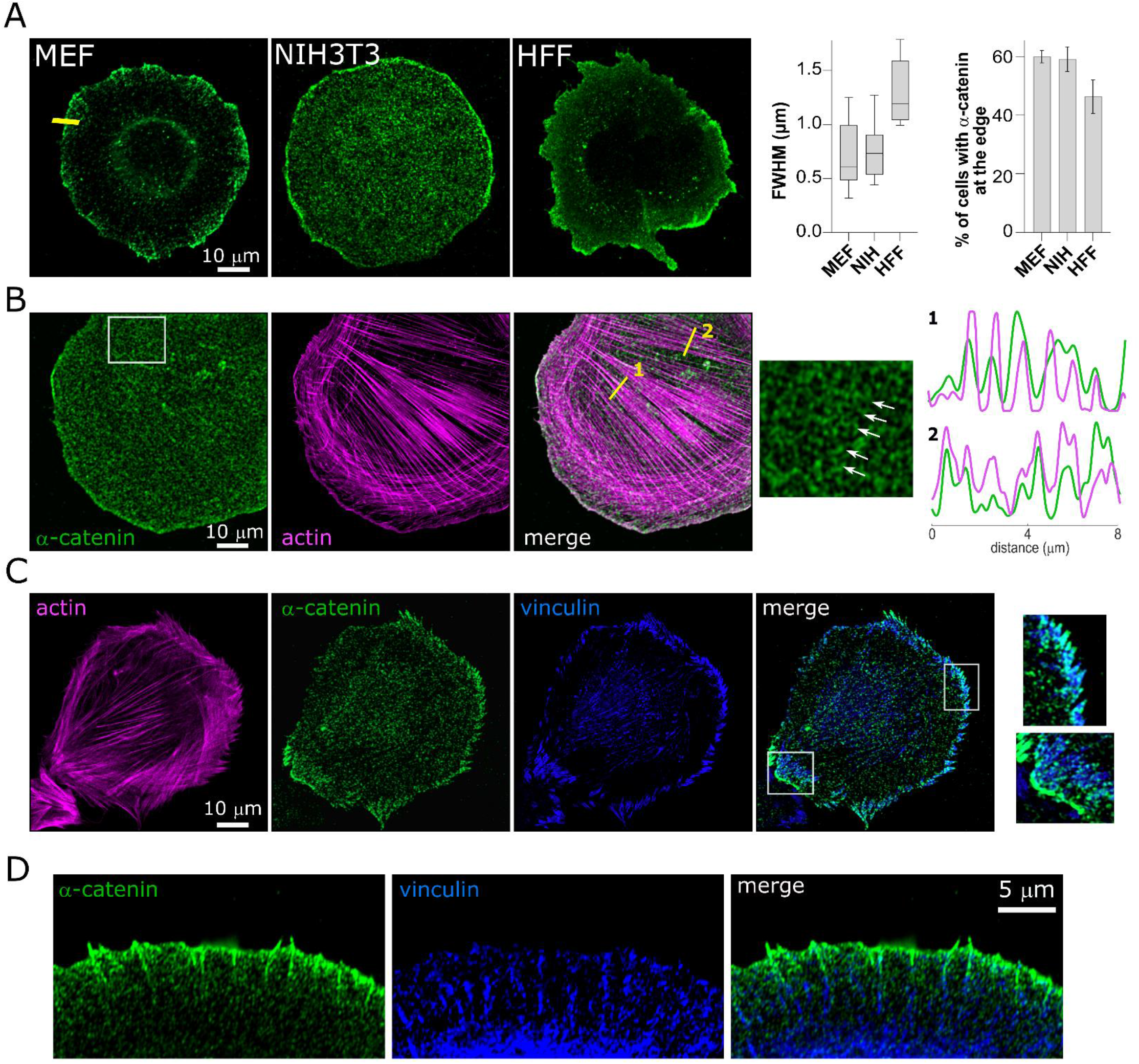
α-catenin localizes in the lamellipodium at early times of cell spreading. (A) Representative images of three fibroblast cell lines immunostained for α-catenin after 15 min of spreading on FN-coated coverslips (left), along with the width of the catenin band at the cell edge, represented by Full Width Half Maximum (center), and the percentage of cells from each fibroblast cell line with catenin at the cell-edge (right) (N=30 cells in each case). (B) WT MEF Co-stained for α-catenin and F-actin (phalloidin); the right panel is a zoom-in of the box in the left image, showing α-catenin stripes that coincide with stress fibers; the graphs on the right are the normalized intensities of phalloidin (magenta) and α-catenin (green) measured along the yellow lines in the merged image. (C) WT MEF Co-stained for actin, α-catenin, and vinculin; right panel images are zoom-ins of the boxes in the merged image, showing vinculin and α-catenin in cell-matrix (top) and cell-cell (bottom) adhesions. (D) Zoom-in on the edge of a WT MEF cell co-stained for α-catenin and vinculin.

Next, we set out to characterize the interaction of α-catenin in mesenchymal cells with its known binding partners in AJs – F-actin, α-actinin, and vinculin – since they are all key players in integrin adhesions. To that end, we used the MEFs, which express high levels of the αE-catenin isoform (Supplementary Fig. 1A). Notably, the recruitment of α-catenin to the cell edge was independent of vinculin and α-actinin, as their depletion from the cells did not affect the localization of α-catenin (Supplementary Fig. 1B). Co-staining the cells for α-catenin and F-actin showed that they were often colocalized at the cell edge at early stages of cell spreading (Fig. 1B). A close examination of cell centers also revealed striped patterns of α-catenin that coincided with actin stress fibers in the majority of analyzed cells (though these patterns were at times obscured by cytoplasmic staining) (Fig. 1B, Supplementary Fig. 1C)

Next, we co-stained the cells for α-catenin, α-actinin, and F-actin. Consistent with previous studies, during early cell spreading, α-actinin appeared in nascent adhesion sites and was also recruited to the cell edges, where it was co-localized with F-actin (Supplementary Fig. 2A) (Meacci et al., 2016; Roca-Cusachs et al., 2013). However, close examination of high-resolution confocal images showed that α-catenin and α-actinin were not colocalized at the cell edges (Supplementary Fig. 2B). Similarly, at later time points these two proteins were not colocalized on actin stress fibers (Supplementary Fig. 2C). Thus, α-actinin and α-catenin overlaid actin filaments in a non-overlapping manner.

**Figure 2.**
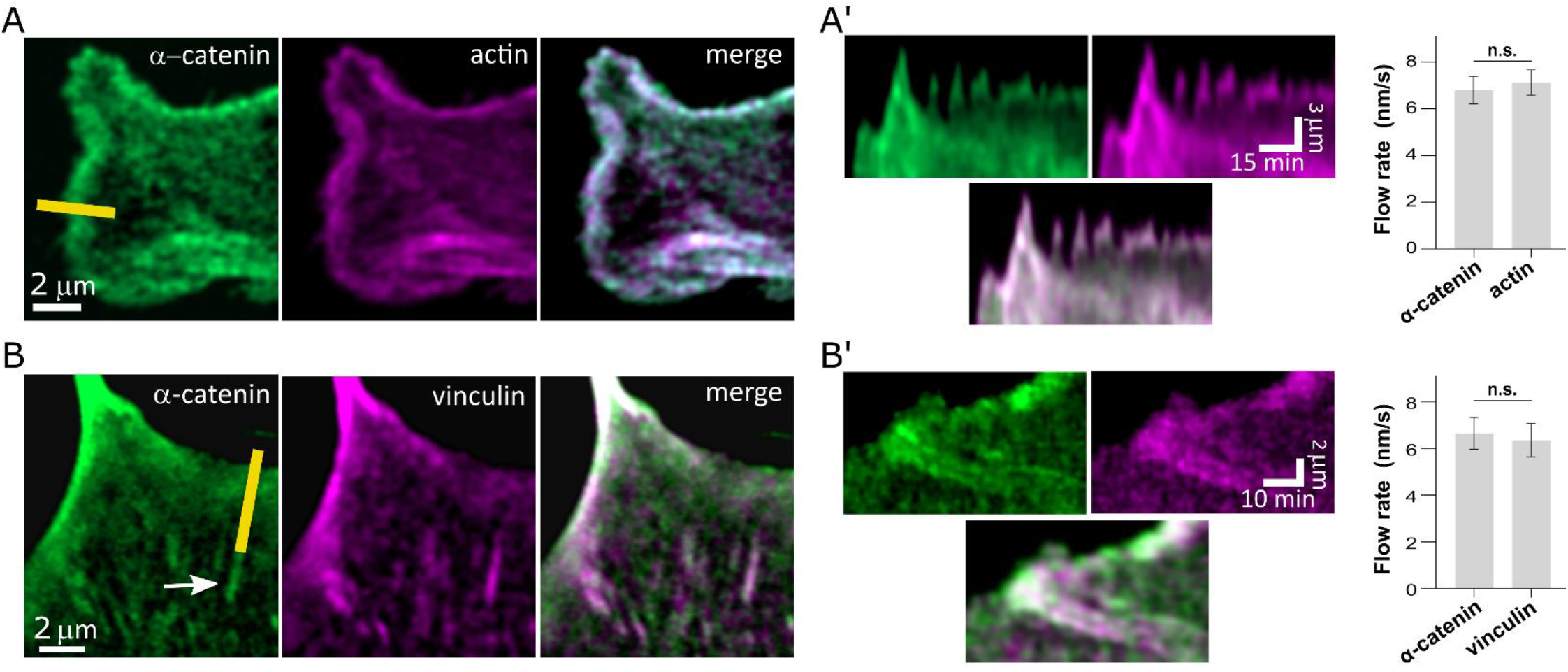
α-catenin flows from the cell edge with actin and vinculin. (A) Frame from a movie of a cell expressing GFP-α-catenin and td-Tomato-Tractin showing colocalization of actin and α-catenin at the edge; (A’) Kymographs taken from the yellow line shown in panel A. The chart on the right shows quantifications of the flow rates of α-catenin and actin in adhesions (N = 18 adhesions in each case). (B) Average of 6 frames (equivalent to 2 min) from a movie of a cell expressing GFP-α-catenin and mCherry-vinculin showing colocalization at the edge as well as in mature adhesions (arrow); (B’) Kymographs taken from the yellow line shown in panel B. The chart on the right shows quantifications of the flow rates of α-catenin and vinculin in adhesions (N = 24 adhesions in each case). Statistical analysis for the flow rates was performed with Student’s t-test followed by Welch’s correction (*, *p* < .05; **, *p* < .01; ***, *p* < .001, ****, *p* < .0001).

We next turned to test the relative distributions of α-catenin and vinculin. In cells that displayed nascent vinculin adhesions at the cell edge, we found similar distributions of α-catenin, though colocalization was partial (Fig. 1C). Also, in some cases, in cells that displayed mature vinculin adhesions 1-3 μm away from the α-catenin-rich cell edges, we observed α-catenin patches extending from the edges towards these adhesions (Fig. 1D), suggesting that α-catenin was flowing from the cell edges toward integrin adhesions and interacting with them. This was observed in a relatively small number of cases, suggesting that such an interaction might be transient and would be better detected by live cell imaging.

### α-catenin undergoes retrograde flow with F-actin and interacts with integrin adhesions

To further explore the relationship of α-catenin with actin, vinculin and α-actinin in the context of the lamellipodium and integrin adhesions, we performed live-cell imaging of MEFs on FN-coated coverslips. Tracking the dynamics of wild-type (WT) GFP-α-catenin along with the F-actin marker tdTomato-Tractin (Belin et al., 2014) showed that the two markers were predominantly colocalized, and displayed the same protrusion-retraction cycles at the cell edge (Fig. 2A-A’, Supplementary Video 1). Moreover, GFP-α-catenin displayed the same flow patterns as that of F-actin from the cell edge inwards, and decorated actin stress fibers as they gradually grew over time (Supplementary Video 1). Consistent with the immunostaining results, imaging live cells co-expressing GFP-α-catenin and mCherry-α-actinin showed that they did not overlap (Supplementary Video 2). In particular, during protrusion-retraction cycles, when α-actinin was associated with the protrusion phase, α-catenin was associated with the retraction phase, and vice versa (Supplementary Fig. 2D-D’). Thus, α-catenin and α-actinin do not interact within the context of cell edge dynamics and integrin adhesions.

In contrast, imaging live cells co-expressing GFP-α-catenin and mCherry-vinculin showed that they had similar dynamics and flow patterns. In particular, α-catenin co-localized with vinculin within adhesions that were growing and sliding from the cell edge toward the center (Fig. 2B’, Supplementary Video 3), suggesting a specific interaction between α-catenin and maturing FAs. Indeed, temporal averaging of GFP-α-catenin videos revealed patterns of α-catenin expression that matched the localization of FAs situated 1-3 μm inwards from the cell edge (Fig. 2B).

### α-catenin interacts with vinculin in focal adhesions

To further explore the interaction between α-catenin and FAs, we next tested whether a specific connection between α-catenin and vinculin was an underlying reason for their similar flow patterns. Indeed, co-immunoprecipitation (co-IP) showed a direct interaction between the two proteins, which was lost upon α-catenin knockdown (KD) and restored upon expression of WT GFP-α-catenin (Supplementary Fig. 3A). We therefore used a variant of GFP-α-catenin containing a lysine-to-proline mutation at site 344 (L344P) which displays drastically diminished binding to vinculin (Peng et al., 2012; Seddiki et al., 2018). We expressed this variant, along with tdTomato-Tractin or mCherry-vinculin, in α-catenin KD MEFs, and performed live-cell tracking of the cells. This revealed stark differences in the α-catenin and adhesion dynamics compared to those detected in WT GFP-α-catenin cells. First, in ∼61% of the analyzed L344P mutant cells (14 out of n = 23 cells), stable adhesions failed to form, and as a result the cell edges did not stabilize following protrusions, leading to extensive ruffling (Fig. 3A, Supplementary Video 4). This was confirmed using live brightfield imaging of early spreading cells in which L344P mutant cells, and α-catenin KD cells, displayed extensive ruffling compared to WT controls (Supplementary Videos 8-11; Supplementary Fig. 3B). Second, in ∼26% of the cells, the cell edges were able to stabilize without the formation of lamellipodia, and instead the cells spread by projecting narrow protrusions (Fig. 3B). However, even though relatively stable vinculin-containing adhesions were present in such protrusions, these adhesions rarely slid towards the cell center, despite the observed retrograde flow of GFP-α-catenin-L344P on top of them (Fig. 3B, Supplementary Video 5). Evidently, the flow rate of GFP-α-catenin-L344P was considerably higher than that of mCherry-vinculin in these adhesions (12.4±1.1 vs. 6.4±0.7 nm/s; n = 20 for each) (Fig. 3B, Supplementary Video 5). Finally, at the cellular-level, GFP-α-catenin-L344P was often located on actin stress fibers that were attached to vinculin adhesions, but unlike cells expressing WT GFP-α-catenin, the stress fibers were highly contractile and dynamic. This led to the aggregation of GFP-α-catenin-L344P and F-actin at the cell center, often in the form of thick actin bundles near the nucleus (Fig. 3C-D, Supplementary Videos 6 & 7), which was observed in ∼75% of the cells. These three behaviors demonstrate that inhibited interactions between α-catenin and vinculin affects both the adhesions and the actin cytoskeleton.

**Figure 3.**
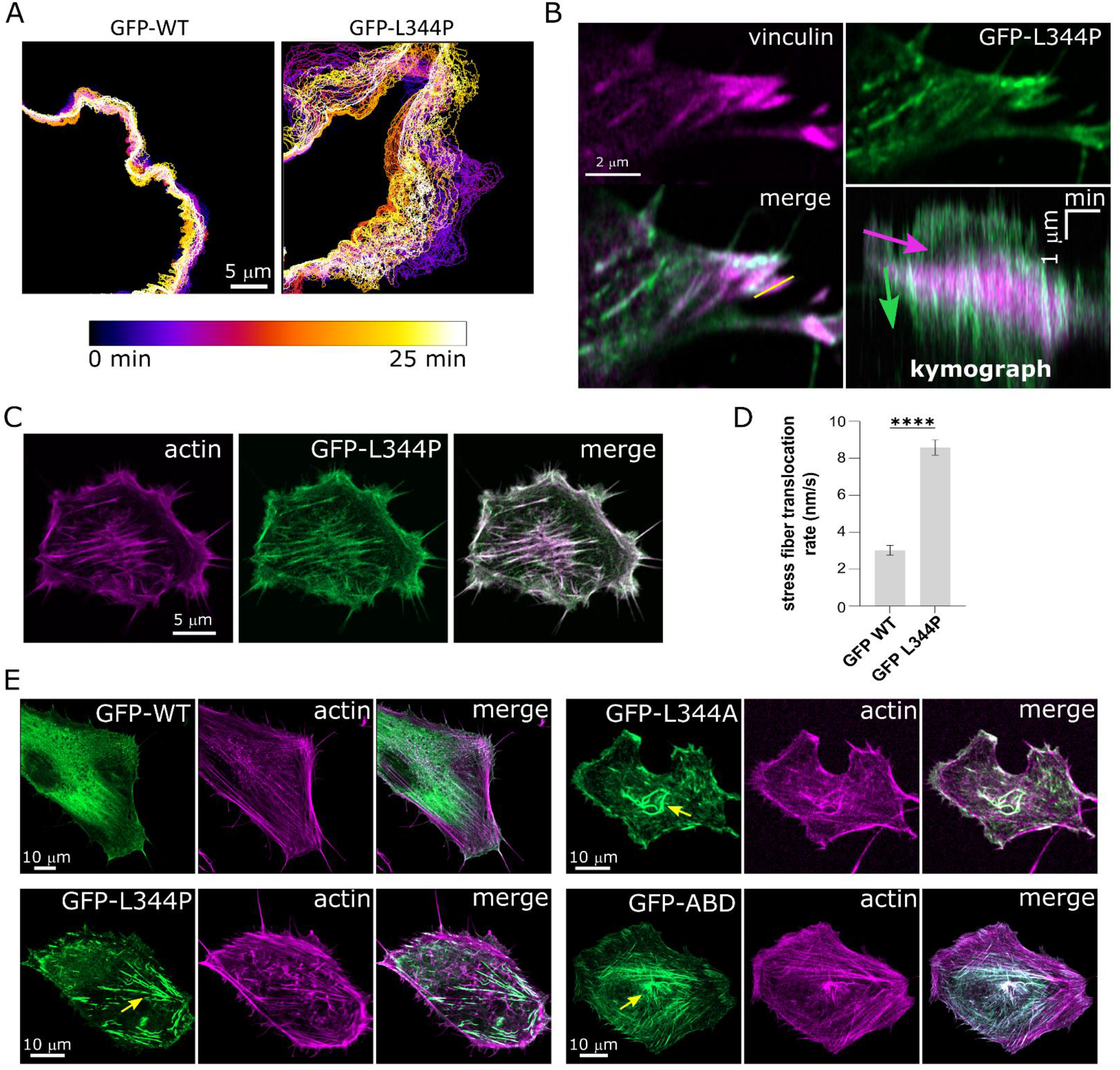
α-catenin-vinculin interaction regulates cell edge activity and stress fiber formation. (A) color-coded time-series of the cell edge of α-catenin KD cells expressing WT GFP-α-catenin (left) and GFP-α-catenin L344P (right), showing much higher cell edge activity in the latter. (B) Average of 6 frames (equivalent to 2 min) from a movie zoomed-in on the edge of a cell expressing GFP-α-catenin L344P and mCherry-vinculin; the bottom right image is a kymograph taken from the yellow line shown in the bottom left image. Note the difference in speed (slope) between vinculin (purple arrow) and α-catenin L344P (green arrow). (C) Frame from a movie (Supplementary Video 6) of an α-catenin KD cell expressing GFP-α-catenin L344P and td-Tomato-Tractin showing aggregation of both at the cell center. (D) Rate of translocation of stress fibers in cells expressing WT GFP-α-catenin and GFP-α-catenin L344P, as measured from kymographs specifically focused on transverse arc type of stress fibers. N>25 stress fiber retraction events from 8 cells each. (E) Micrographs of α-catenin KD cells expressing WT GFP-α-catenin, GFP-α-catenin L344P, GFP-α-catenin L344A, and GFP-ABD, stained for F-actin (phalloidin). The yellow arrows point to actin aggregates near the cell center. Statistical analysis for the stress fiber translocation rate was performed with Student’s t-test with Welch’s correction (*, *p* < .05; **, *p* < .01; ***, *p* < .001, ****, *p* < .0001).

Notably, the enhanced localization of the L344P mutant in regions rich with actin stress fibers suggested that it could bind these structures more efficiently compared to WT α-catenin. To test this, we expressed three α-catenin variants in the cells: WT GFP-α-catenin, GFP-α-catenin-L344P, or a GFP-labeled fragment of α-catenin corresponding to its ABD (amino acids 680-906 (Nicholl et al., 2018)). Furthermore, as proline is known to induce bending within α-helices, we considered that the L344P mutation could potentially disrupt the structure of the second α-helix present in the M_I_ domain of α-catenin (Abe et al., 2013), thereby giving rise to unknown structural changes in other areas of the protein. Hence, to verify that the observed effects of the L344P mutation were not due to α-catenin misfolding, we added a mutant form of GFP-α-catenin in which the lysine at position 344 was replaced by alanine (GFP-α-catenin-L344A). Staining the cells for F-actin showed that all three mutated forms of α-catenin displayed similar localization on stress fibers in the majority (>80%, n = 20 cells in each case) of the transfected cells and in central regions of the cells, in the form of perinuclear actin bundles (Fig. 3E). WT GFP-α-catenin appeared primarily at the cell edges (85% of the cells, n = 20 cells) and on stress fibers, but this was often obscured by cytoplasmic localization (60% of the cells) (Fig. 3E), and the cells rarely (<5%) formed perinuclear actin bundles. Thus, the enhanced contrast between stress fiber and cytoplasmic localization of the three mutant forms, combined with the formation of actin bundles in the presence of these mutants, suggest that “free” α-catenin with an active ABD can bind efficiently to stress fibers, potentially affecting their contractility.

Collectively, these results demonstrate that the α-catenin–vinculin interaction is necessary for normal cell spreading, regular protrusion/retraction cycles of the cell edge, sliding of integrin adhesions, and proper formation of actin stress fibers.

### α-catenin affects force transmission to the matrix

The results described above indicate that vinculin may act as a clutch that engages with α-catenin within integrin adhesions as the latter is bound to actin fibers. Indeed, introducing the L344P mutation in α-catenin KD MEFs caused a noticeable disconnect between the respective flows of α-catenin and vinculin (compare Fig. 2B and Fig. 3B). Quantification of the flow rates within adhesions showed that the α-catenin–vinculin interaction attenuated the flow of α-catenin by ∼2-fold (12.4±1.1 and 6.3±0.6 nm/s for L344P and WT α-catenin, respectively; n = 15 each), thus reinforcing the idea that the α-catenin–vinculin interaction acts as a clutch. Therefore, we next set out to test the effect of this interaction on force transmission to the matrix, since stronger integrin-actin engagement leads to more efficient force transmission (Elosegui-Artola et al., 2016). To that end, we plated the cells on arrays of FN-coated pillars and measured the time-dependent deformation of the pillars (Feld et al., 2020). As shown in Fig. 4A, WT MEFs gradually displaced the pillars over a period of about 10 min, until reaching a plateau of approximately 120 nm. In agreement with the adhesion maturation results, α-catenin KD led to a significant decrease in matrix deformation (maximal displacement of ∼40 nm). Furthermore, WT GFP-α-catenin expression restored pillar displacement almost completely (maximal displacement of ∼100 nm) whereas GFP-α-catenin-L344P expression did not (Fig. 4A). These results strengthen the notion that α-catenin–vinculin binding is a crucial clutch element within adhesions that is required for proper contractile activity and force transmission into the matrix.

**Figure 4.**
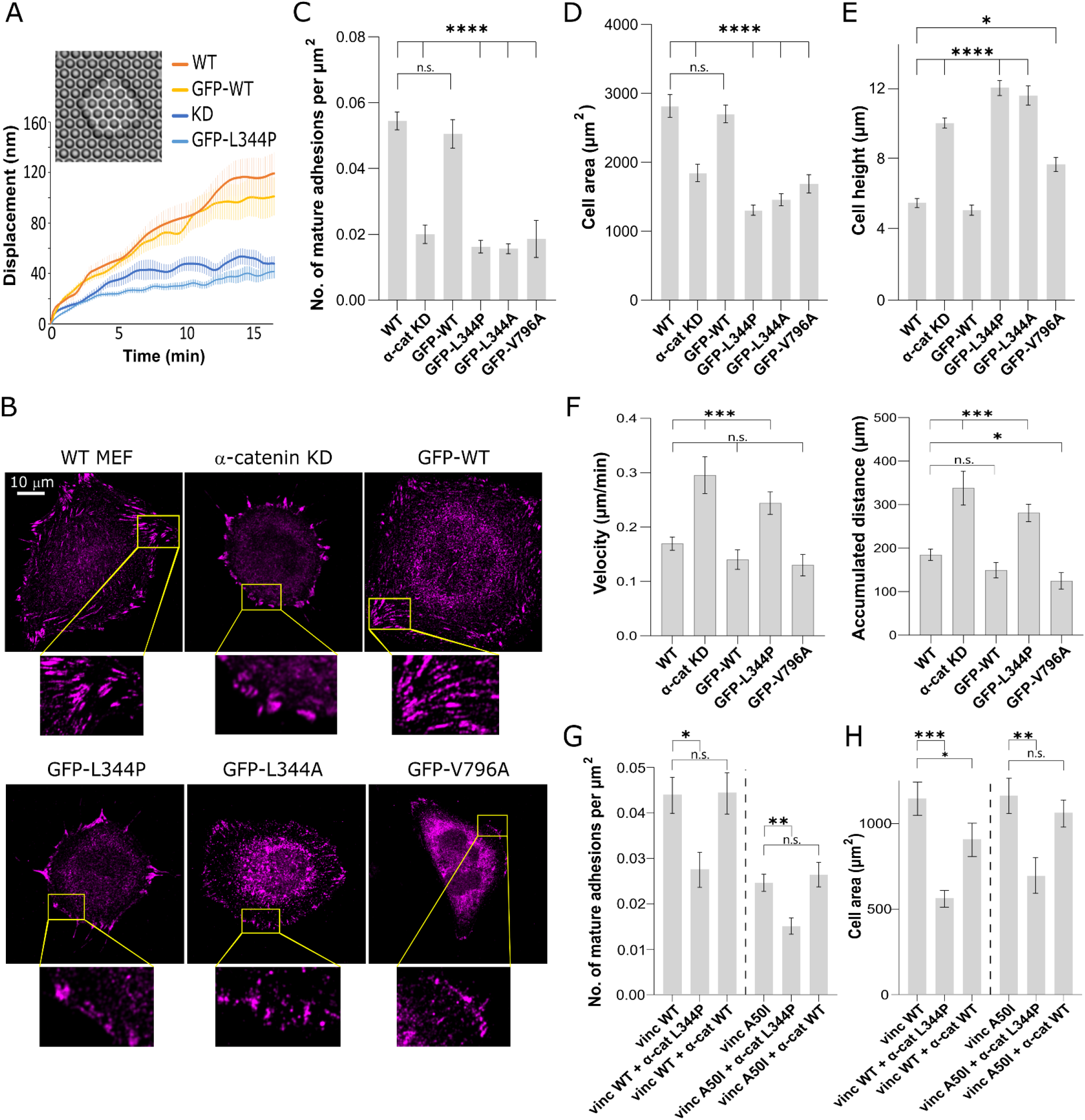
α-catenin L344P affects force transmission and adhesion maturation. (A) Displacement as a function of time of 2μm diameter FN-coated pillars by MEFs spreading on top of them (see example in inset image). The graphs shown are mean ± SEM (in lighter hues). N>30 pillars in each case. (B) Micrographs of a WT MEF, α-catenin KD MEF, and α-catenin KD MEFs expressing WT GFP-α-catenin / GFP-α-catenin L344P / GFP-α-catenin L344A / GFP-α-catenin V796A, all immunostained for vinculin after 6 hours of spreading on FN-coated coverslips (GFP channel not shown). (C-E) quantifications of the number of mature adhesions per μm^2^, cell areas, and cell heights of the six cell types 6 hours after plating on FN-coated coverslips (N>20 cells in all cases). (F) Velocities and accumulated distances measured from MEF single cell motility videos (N>17 cells in all cases). (G-H) quantifications of the number of mature adhesions per μm^2^ and areas of vinculin-/-MEFs expressing WT mCherry-vinculin or mCherry-vinculin A50I, alone or together with WT GFP-α-catenin or GFP-α-catenin L344P (N>15 cells in all cases). Statistical analysis of the adhesions, cell areas, cell heights and cell motility were tested by two-way ANOVA followed by Tukey’s multiple comparisons test (*, *p* < .05; **, *p* < .01; ***, *p* < .001; ****, *p* < .0001).

### The α-catenin–vinculin interaction is essential for focal adhesion maturation

Next, we sought to test the effect of α-catenin–vinculin interactions on the growth of nascent adhesions into mature FAs, as this process is force-dependent (Galbraith et al., 2002; Riveline et al., 2001). To that end, we stained the cells for vinculin and F-actin, and found that whereas WT MEFs formed mature vinculin-containing FAs, α-catenin KD cells mostly did not display mature vinculin adhesions, and had almost exclusively small adhesions (‘focal complexes’) at the cell edge (Fig. 4B-C). In line with the reduction in mature adhesions, α-catenin KD cells failed to spread out and flatten, as evident by their reduced area (Fig. 4D) and increased height (Fig. 4E) compared to WT cells. Similarly, knocking down α-catenin in NIH3T3 cells resulted in the formation of small adhesions and a decrease in cell spreading (Supplementary Fig. 4).

Expression of GFP-α-catenin in the α-catenin KD MEFs restored the formation of mature FAs, but, importantly, KD cells expressing GFP-α-catenin-L344P or L344A displayed similar adhesions as those of α-catenin KD cells (Fig. 4B-C). Similar results were obtained when staining for zyxin (Supplementary Fig. 5A), a marker for mature FAs (Zaidel-Bar et al., 2003), confirming that the observed lack of mature vinculin-containing adhesions was not due to lack of vinculin recruitment into mature adhesions but rather due to the absence of such adhesions all together. Furthermore, α-catenin KD cells and cells expressing GFP-α-catenin-L344P or L344A were significantly smaller (Fig. 4D) and less flat (Fig. 4E) compared to WT cells or cells expressing WT GFP-α-catenin, consistent with their inability to form mature adhesions.

**Figure 5.**
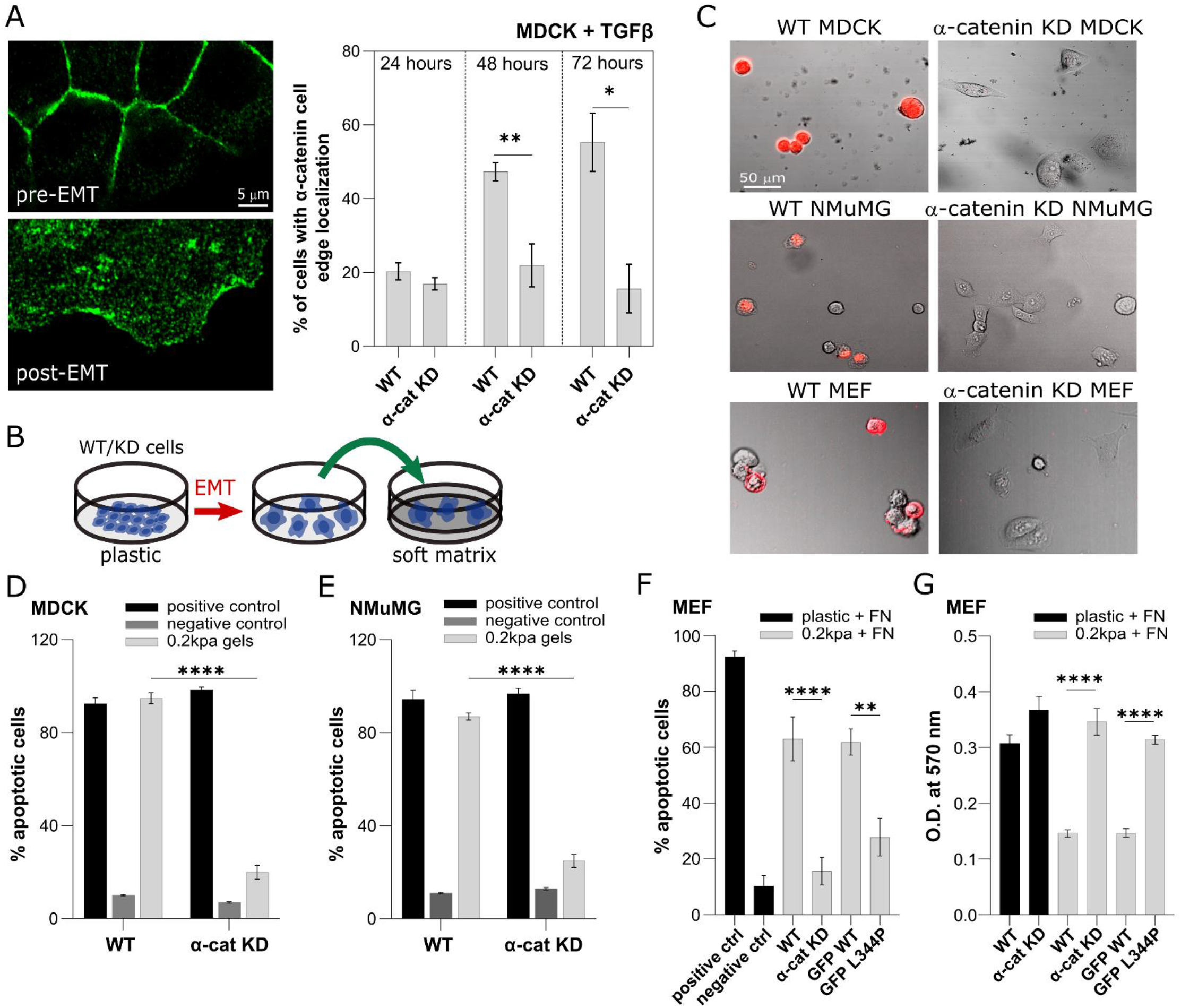
α-catenin does not affect rigidity-dependent EMT. (A) Left: α-catenin staining in WT MDCK cells before and after EMT. Right: quantification of the percentage of cells with α-catenin cell edge localization, 24, 48 and 72h after the addition of TGF-β. (B) Schematic of the experiment designed to test α-catenin’s role in rigidity-dependence post-EMT in MDCK and NMuMG cells. (C) Representative images of MDCK, NMuMG (after EMT) and MEF cells stained with Annexin-FITC on 0.2kPa gels coated with FN, 6 hours after plating (PI staining not shown). (D) Quantification of the percent of Annexin V & PI positive WT and α-catenin KD MDCK cells on 0.2kPa gels coated with FN. (E) Quantification of the percent of Annexin V & PI positive WT and α-catenin KD NMuMG cells on 0.2kPa gels coated with FN. (F) Quantification of the percent of Annexin V & PI positive WT MEFs, α-catenin KD MEFs, and α-catenin KD MEFs expressing WT GFP-α-catenin or GFP-α-catenin L344P plated for 24 hours on 0.2kPa FN-coated matrices. (G) MTT measurement of WT MEFs, α-catenin KD MEFs, and α-catenin KD MEFs expressing WT GFP-α-catenin or GFP-α-catenin L344P plated for 24 hours on 0.2kPa FN-coated matrices. In the Annexin/PI graphs, positive and negative controls, the cells were heated at 65°C for 15 minutes for the former and left untreated for the latter and plated on FN-coated plastic ibidi wells. For the MTT assay, the positive growth control refers to WT and α-catenin KD MEF cells plated on a FN-coated 96-well plate (plastic). Statistical analysis of the α-catenin cell-edge localization, %of apoptotic cells and MTT assay were tested by two-way ANOVA followed by Tukey’s multiple comparisons test (*, *p* < 05; **, *p* < .01; ***, *p* < .001; ****, *p* < .0001).

Next, we expressed an α-catenin variant in which valine 796 – a key residue within the ABD – was replaced by alanine (V796A), as this mutant is known to reduce the binding ability of α-catenin to actin (Ishiyama et al., 2018). Cells expressing this mutant showed similar results (Fig. 4B-E) to the α-catenin KD cells and the other mutants, suggesting that the actin-binding ability of α-catenin is important to adhesion maturation as well. Interestingly, single cell motility assays showed that α-catenin KD cells and cells expressing the L344P mutant were more motile compared to the WT controls, as evident by their increased distances and velocities (Fig. 4F). In contrast, V796A mutant cells displayed a similar motility phenotype to that of WT controls (Fig. 4F), despite displaying adhesion and cell size phenotypes comparable to the KD and L344P cells (Fig. 4C-E). This disengagement of phenotypes points to a complex relationship between the strength of α-catenin–F-actin binding, adhesion reinforcement, and cell migration, which should be investigated in more depth in future studies.

As a major axis of vinculin recruitment to FAs is through its binding to talin, we next wished to test the effects of this interaction on adhesion growth in the context of the α-catenin–vinculin interaction. To that end, we transfected vinculin-/-fibroblasts with WT mCherry-vinculin or mCherry-vinculin-A50I – a mutant form of vinculin which displays reduced binding to talin compared to WT vinculin, but is still recruited to adhesions (Cohen et al., 2006). We co-expressed WT GFP-α-catenin or GFP-α-catenin-L344P constructs in these cells. As shown in Fig. 4G, despite the fact that the A50I mutation led to formation of less mature adhesions compared to WT vinculin (as expected), co-expression with the L344P mutant led to a further decrease in adhesion sizes, which was accompanied by a reduction in cell area (Fig. 4H). These results indicate that once vinculin is recruited to adhesions (either through talin or through other adhesion proteins), its interaction with α-catenin is essential for the regulation of adhesion maturation and cell spreading.

To further test the relevance of the α-catenin–vinculin interaction in the regulation of integrin adhesions, we used Matrigel (basement membrane-like) matrices, which were recently shown to induce the sliding of adhesions from the cell edge towards the center, resulting in highly elongated adhesions around the nucleus (Lu et al., 2020). We plated the cells on Matrigel-coated coverslips, and stained them for vinculin, which revealed that WT MEFs and WT GFP-α-catenin MEFs formed a significantly higher number of sliding adhesions than the α-catenin KD cells and cells expressing GFP-α-catenin-L344P (Supplementary Fig. 5B).

Taken together, these results show that the α-catenin–vinculin interaction plays a critical role in integrin adhesion regulation. This raises the question of whether such a role could be affected by the presence of cadherin-based cell-cell adhesions in the cells. To address this, we studied MEFs which were in contact with one another (mesenchymal cell-cell) as well as with the matrix. We observed the presence of α-catenin, β-catenin, and N-Cadherin in the mesenchymal cell-cell contacts, and found that β-catenin (but not N-Cadherin) was also present at the lamellipodia along with α-catenin (Supplementary Fig. 5C). Still, in such cells, mature vinculin-containing FAs formed in lamellipodial regions that included α-catenin (Supplementary Fig. 5C), suggesting that α-catenin can affect both cell-cell and cell-matrix adhesion regulatory processes within the same cell.

### α-catenin regulates rigidity-dependent cell growth following EMT

Since the α-catenin–vinculin complex has a mechanosensory role within E-cadherin adhesions (Thomas et al., 2013), we aimed to test whether it plays a similar role in ECM rigidity mechanosensing by integrin adhesions. Indeed, the observed decrease in matrix deformation upon the loss of α-catenin–vinculin binding (Fig. 4A) suggests that this interaction is required for the proper transmission of contractile displacements involved in mechanosensing (Feld et al., 2020). We first focused on EMT, which was shown to occur on stiff matrices but not on soft ones (Wei et al., 2015), and set out to test whether α-catenin’s dual role in E-cadherin adhesions and in integrin adhesions might be critical for this phenomenon. Therefore, we used the epithelial Madin-Darby Canine Kidney (MDCK) cell line, as well as an MDCK variant in which α-catenin expression was knocked-down (Benjamin et al., 2010) (Supplementary Fig. 6A). First, to test the effect of α-catenin KD, we grew the cells for 48h on FN-coated glass coverslips and quantified the expressions of E-cadherin, vimentin, and αSMA. WT MDCKs displayed a strong epithelial phenotype with tight E-cadherin connections between the cells and very low vimentin levels (Supplementary Fig. 6B). α-catenin KD MDCKs were less tightly packed compared to the WT cells, and displayed E-cadherin adhesions, which was accompanied with a slight increase in vimentin and α-SMA intensity (Supplementary Fig. 6B-C). These results indicated that the loss of α-catenin was not sufficient for the cells to undergo complete EMT. We next tested if the absence of α-catenin could affect EMT occurrence as a function of matrix rigidity. To that end, we plated both cell lines on soft (0.2 kPa) and stiff (25kPa) FN-coated substrates for 3 hours before inducing EMT by adding TGFβ. After a 72h incubation, WT cells on the soft matrix remained epithelial (Supplementary Fig. 6D), whereas on the stiff matrix, the cells formed colonies that were less tightly packed, with cells that were detached from the colonies at the periphery (Supplementary Fig. 6D, yellow arrows), indicative of partial EMT (Supplementary Fig. 6D). Interestingly, on soft matrices the KD cells also displayed a partial EMT phenotype, while on stiff matrices they underwent complete EMT, as evident by the complete loss of cell-cell contacts and spindle-like morphologies, which are characteristic of mesenchymal cells (Supplementary Fig. 6D). Thus, although the baseline phenotypes of WT and α-catenin KD MDCK cells on soft matrices differed in the presence of TGFβ (fully epithelial and partial EMT, respectively), stiff matrices shifted both cell types toward the mesenchymal state (partial EMT and complete EMT, respectively).

While the EMT assay results suggest that high stiffness can promote EMT in the absence of α-catenin, they provide no indication regarding the role of α-catenin post-EMT. Notably, we found that during EMT induction, WT MDCK cells displayed a gradual increase in the cell edge localization of α-catenin (Fig. 5A). We therefore wished to explore whether α-catenin had a rigidity-dependent role in the mesenchymal state, post-EMT. To that end, we induced EMT of WT and α-catenin KD MDCK cells in plastic dishes and then transferred them onto soft (0.2 kPa) FN-coated substrates (Fig. 5B). Remarkably, Annexin V & PI staining showed that only ∼20% of α-catenin KD MDCK cells activated apoptosis 6 hours after plating, compared to ∼95% of WT MDCK cells (Fig. 5C,D). We observed similar results with another epithelial cell line, NMuMG, in which we knocked-down the expression of α-catenin (Fig. 5C,E, Supplementary Fig. 6A). Furthermore, in the MEFs, α-catenin KD and the L344P mutation both led to apoptosis inhibition on 0.2 kPa FN-coated surfaces, compared to WT controls (Fig. 5C,F). Importantly, using the MTT assay we verified that while WT MEFs and MEFs expressing WT GFP-α-catenin failed to grow on the 0.2 kPa surfaces, α-catenin KD and L344P mutant cells were able to proliferate on such matrices (Fig. 5G, Supplementary Fig. 6E).

Collectively, these results show that the ability of α-catenin to interact with vinculin is critical for cellular mechanosensing of ECM rigidity, and regulates the anchorage-dependent growth of cells in the mesenchymal state.

## Discussion

In this study, we identify a critical role for α-catenin as a regulator of integrin adhesions and the associated actin cytoskeleton. We show that α-catenin is recruited to lamellipodial regions, where it interacts with actin that is undergoing retrograde flow toward the cell center (Figs. 1&2). As actin travels past integrin adhesions along with α-catenin, the latter interacts directly with vinculin in the adhesions (Fig. 3). This interaction is important for force transmission from the actin cytoskeleton through the adhesions, and for the maturation of focal complexes into FAs (Fig. 4), which is consistent with the concept of adhesion growth being a force-induced process. At later stages, α-catenin decorates stress fibers in regions devoid of α-actinin, and the L344P mutant (which has diminished α-catenin–vinculin binding) leads stress fibers to become highly contractile and aggregated at the cell center, suggesting that the α-catenin–vinculin link affects stress fiber contractility. The underlying mechanism of this phenomenon is not clear and should be addressed in future studies. One possibility is that the mere presence of α-catenin enhances stress fiber contractility; hence, the sequestration of α-catenin by vinculin in the adhesions could prevent excess α-catenin translocation into the stress fibers, thereby attenuating excess contractility. Another possibility is that the poor connection between stress fibers and FAs in the presence of the L344P mutant limits the transmission of the contractile forces to the ECM, thereby leading to enhanced deformation of the stress fibers themselves, as predicted by our recent study (Feld et al., 2020).

The classical model for integrin adhesion maturation involves the talin-vinculin module, and is described as a force-dependent mechanism in which talin stretching reveals hidden vinculin binding sites (del Rio et al., 2009). Thus, upon vinculin binding, the link between the cytoplasmic tails of the integrins and the actin cytoskeleton is reinforced, thereby stabilizing the adhesions (Elosegui-Artola et al., 2016). Although the relevance of this model to the behavior of cells on soft and stiff matrices is still being debated (Driscoll et al., 2020), the talin–vinculin module is clearly important for adhesion regulation and mechanosensing of the ECM. While vinculin has an ABD, it appears that its ability to bind to α-catenin is equally vital for proper interaction with F-actin, and consequently for proper force transmission and adhesion maturation. Notably, the α-catenin–vinculin interaction does not depend on vinculin–talin binding, as vinculin A50I behaved similarly to WT vinculin (Fig. 4G-H). The results we report here on the role of α-catenin thus provide an important insight on the link between vinculin and actin, and the classical adhesion maturation model.

Normally, cells can detect the rigidity of the extracellular environment by assembling the mechanosensing machinery, which includes components such as talin, vinculin, α-actinin, and tropomyosin 2.1 (Wolfenson et al., 2019). When cells are on stiff matrices, they form large and mature adhesions that promote their proliferation, but on extremely soft matrices (e.g., 0.2 kPa) the adhesions fail to mature, and the cells undergo anoikis. Here, we show that α-catenin is an essential component of this machinery, and the absence of α-catenin–vinculin interactions leads to impaired mechanosensing, which manifests in the ability of cells to grow on soft matrices (anchorage-independent conditions) (Fig. 5). This observation is in line with our previous findings which show that anchorage-independent cells often display smaller adhesions and spread less on stiff matrices compared to soft matrices (Wolfenson et al., 2016; Feld et al., 2020). We hypothesize that this behavior could stem from an altered balance between cellular force production and adhesion reinforcement, and we aim to test this in future studies.

Interestingly, we find that the loss of α-catenin is not necessary for the cells to undergo EMT, however, the role of α-catenin in the process of rigidity-dependent EMT is not to be disregarded. In particular, our results show that the epithelial MDCK cell line undergoes (partial) EMT on soft surfaces upon treatment with TGFβ only when α-catenin is absent from the cells. Strikingly, we found that the most significant role of a-catenin lies in the post-EMT stages. The absence of α-catenin endows cells with the ability to grow on soft matrices in the mesenchymal state after EMT, which we observed in MDCK and NMuMG cells, as well as in the mesenchymal MEFs. The tumor suppressive role of α-catenin is typically attributed to its role in the maintenance of AJs (Bajpai et al., 2009) and/or its involvement in cytoplasmic sequestration of pro-proliferative transcriptional regulators (Piao et al., 2014; Silvis et al., 2011). Our findings add an additional layer to this picture, as they indicate that α-catenin also plays a significant role in regulating integrin adhesions and mechanosensing, which is directly linked to anchorage-independence (Wolfenson et al., 2016; Meacci et al., 2016; Yang et al., 2019), a hallmark of cancer cells (Guadamillas et al., 2011).

## Materials and Methods

### Cell culture, reagents, and transfections

WT MEFs RPTPα^+/+^ cells (henceforth known as MEFs), NIH3T3, WT MDCK, and MDCK α-catenin KD were received from of MBI Singapore. MEF vinculin^-/-^ cells were received from Benny Geiger’s lab (Weizmann Institute of Science). NMuMG cells were received from Yaron Antebi’s lab (Weizmann Institute of Science). All cells were cultured at 37° C in a 5% CO2 incubator in Dulbecco’s Modified Eagle Medium (DMEM) supplemented with 10% fetal bovine serum, and 100 IU/ml Penicillin-Streptomycin (all reagents were from Biological Industries). Recombinant TGFβ (10 ng/ml) were purchased from Peprotech (Rocky Hill, NJ). For EMT experiments, the cells were treated with TGFβ for 48-72 hours.

Transfections were carried out 1 day before measurements using the NEPA21 Electroporator (Nepa Gene) according to the manufacturer’s instructions, with ∼10^6^ cells per reaction and 10 μg DNA.

### Plasmids and shRNA oligonucleotides

The plasmids for GFP/mCherry-tagged α-catenin, α-actinin, and vinculin, as well as the Tomato-Tractin plasmid, were obtained from MBI Singapore. The L344P, L344A, V796A mutations were inserted into the GFP-α-catenin plasmid using the Q5 Site-Directed Mutagenesis Kit (New England Biolabs).

Lentiviral KD of α-catenin was performed using the SHCLNG-NM_009818 MISSION® shRNA plasmid (Merck); control cells were generated using the SHC202 -MISSION® TRC2 pLKO.5-puro Non-Mammalian shRNA Control Plasmid (Merck). After infection, cells were grown in 1 and 4 μg/ml puromycin for the MDCK and MEF, and KD was tested using Western blotting and immunofluorescence measurements.

### Pillars, soft gel fabrication, and cell spreading

Pillar fabrication was done by pouring PDMS (Sylgard 184, Dow Corning; mixing ratio – 10:1) into silicon molds (fabricated as previously described (Ghassemi et al., 2012)) with holes at fixed depths and distances. The molds were then placed, face down, onto glass-bottom 35 mm dishes (#0 coverslip, Cellvis) which were incubated at 65°C for 12h to cure the PDMS. The molds were next peeled off while immersed in ethanol to prevent pillar collapse, which was then replaced by serial dilutions with PBS. Human plasma full-length fibronectin (Merck) was added to the dish at a final concentration of 10 μg/μl for a 1h incubation at 37°C. Next, residual fibronectin was washed away by replacing the buffer to HBSS buffer (Biological Industries) supplemented with 20 mM HEPES (pH 7.2) or PBS.

All pillars had a diameter of 2 μm, and heights of 9.4 or 13.2 μm. We used 2 μm diameter pillars as these can be used to measure the long-term time-dependent forces that are generated after initial formation and reinforcement of the adhesions (Wolfenson et al., 2016; Ghassemi et al., 2012). The center-to-center spacing between pillars was 4 μm. Pillar bending stiffness, *k*_*ECM*_, was calculated by Euler–Bernoulli beam theory:

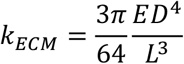

where D and L are the diameter and length of the pillar, respectively, and E is the Young’s modulus of the material (=2 MPa for the PDMS used here).

The 0.2 and 25kPa substrates were fabricated by using Sylgard 52-276, at a ratio of 2:1 and 1:2.7, respectively, according to the measurements performed previously (Ou et al., 2016).

### Pillar displacement measurements

One day prior to the pillar experiments, cells were sparsely plated to minimize cell-cell interactions prior to re-plating. The following day, cells were trypsinized, centrifuged with growth medium, and then resuspended and pre-incubated in HBSS/Hepes at 37°C for 30 min prior to the experiment. Cells were then spread on the fibronectin-coated pillars. In all cases, we made sure that the cells were isolated when plated on the substrates.

Time-lapse imaging of cells spreading on the pillars was performed using an inverted microscope (Leica DMIRE2) at 37°C using a 63x 1.4 NA oil immersion objective. Brightfield images were recorded every 10 seconds with a Retiga EXi Fast 1394 CCD camera (QImaging). The microscope and camera were controlled by Micromanager software (Edelstein et al., 2010). For each cell, a movie of 1-3 hours was recorded. To minimize photo-damage to the cells, a 600 nm longpass filter was inserted into the illumination path.

Tracking of pillar movements over time was performed with ImageJ (National Institutes of Health) using the Nanotracking plugin, as described previously (Wolfenson et al., 2016). In short, the cross-correlation between the pillar image in every frame of the movie and an image of the same pillar from the first frame of the movie was calculated, and the relative x-and y-position of the pillar in every frame of the movie was obtained. To consider only movements of pillar from their zero-position, we only analyzed pillars that at the start of the movie were not in contact with the cell and that during the movie the cell edge reached to them. Drift correction was performed using data from pillars far from any cell in each movie. For each pillar, the displacement curve was generated by Matlab (MathWorks).

### Immunoblotting, immunoprecipitation and immunofluorescence microscopy

For immunoblots, ice-cold Phosphate Buffered Saline (PBS) was used to wash the cells. The cells were lysed using RIPA buffer (Tris-HCl 10 mM, 1% SDS, 10 mg/ml deoxycholate, 150 mM NaCl, 1% NP40 and protease inhibitors cocktail [Roche, Mannheim, Germany]). The total protein samples were separated using by 12% SDS-PAGE and transferred onto nitrocellulose membranes (BioRad). The membranes were blocked using 5% milk and incubated with primary antibodies for α-catenin (1:1000, Santa Cruz, sc-9988) overnight at 4°C. GAPDH (1:10000, Abcam, #ab8245) was used as a loading control. Following this, the membranes were exposed to peroxidase-conjugated goat anti-mouse IgG (1:3000) for 1 hour at RT. EZ-ECL Enhanced Chemiluminescence Detection Kit (Biological Industries, Israel) was used to visualize the protein bands.

For immunoprecipitation, the cell lysates were incubated with 3μg of anti-vinculin (Thermo Fisher Scientific, #700062) and 3μg of anti-α-catenin (Santa Cruz, sc-9988) in immunoprecipitation buffer (20 mM Tris-HCl [pH 7], 0.3 M NaCl, 2 mM EDTA, 1% NP-40) containing 0.2 mM AEBSF Hydrochloride [Calbiochem, #101500]) overnight at 4°C and subsequently rotated with Pierce™ Protein G Magnetic Beads (Thermo Fisher, #88848) for 1h at 4 °C. Protein complexes were washed three times in PBS and subsequently extracted with 1× SDS loading buffer for 3 min at 95 °C. SDS-PAGE, WB, and ECL analyses were performed as discussed above.

For immunofluorescence microscopy, cells were plated on fibronectin-coated coverslips or Matrigel-coated ibidi 8 well chamber slides, fixed with 4% paraformaldehyde (Sigma, #47608) supplemented with 0.2% Triton X-100 (Sigma, #T8787) in PBS. Blocking was performed using 3% Bovine Serum Albumin (Sigma, #A4503) and 0.2% Triton X-100in PBS for 1 hour at RT. Immunolabeling was performed with primary antibodies against α-catenin (1:300, Santa Cruz, sc-9988), vinculin (1:500, Thermo Fisher Scientific, #700062), E-cadherin (1:300, Merck, #U3254), α-actinin (1:500, Abcam, #ab108198), zyxin (1:100, received from Benny Geiger’s lab), vimentin (1:500, Abcam, #ab24525), N-cadherin (1:200, Abcam, #ab18203), β-catenin (1:200, Sigma, #C2206) and α-SMA (1:200, Cell Marque, #A2547) overnight at 4°C. The cells were washed thrice with PBS followed by the addition of Alexa Fluor 405, Alexa Fluor 488, Alexa Fluor 555, or Alexa Fluor 647-conjugated secondary antibodies for 1 hour at RT in the dark. Images were taken with a Zeiss LSM800 confocal microscope using a 20x 0.9NA air objective or 63x 1.4NA oil objective.

### Image analyses and quantifications

Analyses of the cell area and number of mature and sliding adhesions were performed using a home-built Fiji macro (NIH, Bethesda, MD, USA). The confocal images were subjected to an intensity threshold set to select the cell area using the phalloidin channel, and the ‘analyze particles’ tool was used to measure the area of the cell. To account for mature adhesions, a cell mask was generated which measured the cell area from a distance of 2 μm from the cell edge on all sides. The number of adhesions in the cell-center was calculated using the ‘analyze particle’ tool after thresholding the vinculin channel.

### EMT experiments

The MDCK WT, MDCK α-catenin KD, NMuMG WT, and α-catenin KD were sparsely plated on fibronectin coated 0.2 and 25 kPa gels. 3 hours after plating, EGF (50 ng/ml) or TGFβ (10 ng/ml) were added to the media and left for 48-72 hours before fixation and staining. In another set of experiments, the cells were made to undergo EMT on plastic and then plated on the fibronectin coated 0.2kPa PDMS gels.

### Cell Viability assays

The cell viability of the MEFs were assessed using the MTT assay (Merck, #M5655). 1 × 10^4^ cells were seeded on fibronectin-coated plastic and 0.2kPa gels in 96-well plates and incubated in DMEM. After 24h, the MTT reagent was added to the cells at a final concentration of 0.5mg/ml and incubated at 37°C for 3h. The DMEM was removed and 10% SDS in 0.1M HCl was added to dissolve the crystals. The amount of MTT formazan product was measured at 570nm using a microplate reader,

### Apoptosis assays

The apoptosis assay was performed using the Annexin V-FITC Apoptosis Staining/Detection Kit (Abcam, #14085). 1 ×10^4^ cells were seeded on fibronectin coated plastic and 0.2kPa gels on ibidi 8 well chamber slides. For the positive and negative controls, the cells were heated at 65°C for 15 minutes for the former and left untreated for the latter and plated on fibronectin-coated plastic ibidi wells. 6h after plating, 10μl of FITC-Annexin V and 10μl of Propidium Iodide were added to the cells for 15 minutes and incubated in the dark. The stainings of the apoptotic cells were assessed using fluorescence microscopy.

### Single-cell motility assay

The cells were incubated with 400nM of SiR-DNA (Spirochrome, #SC007) in full DMEM, overnight at 37°C in a 5% CO2 incubator. The cells were then trypsinized, centrifuged and re-suspended in colorless DMEM. 1× 10^4^ cells were plated in each well of a fibronectin coated glass-bottom 24 well plates (Cellvis, #P24-0-N) and incubated at 37°C for 1h. The cells where then imaged at 10X magnification every 15 minutes at 37°C and 5% CO2 using an ImageXpress micro system (Molecular Devices). Quantifications were performed using the Manual Tracking plugin in FIJI and the Chemotaxis and Migration Tool (ibidi, downloaded from: https://ibidi.com/chemotaxis-analysis/171-chemotaxis-and-migration-tool.html).

### RNA extraction and quantitative real-time PCR (qRT-PCR)

RNA was extracted from cultured cells using PureLink RNA Mini Kit (12183018A, Invitrogen) and reverse transcription was performed with qScript cDNA Synthesis Kit (95047-100, Quanta Biosciences). In qRT-PCR experiments 10 ng cDNA were used and PCR products were detected using Fast SYBR™ Green Master Mix (4385614, Applied Biosystems). Expression results were normalized to *Gapdh*, and to the indicated control groups (RQ = 2^-ΔΔCt^). The primers used are listed below:

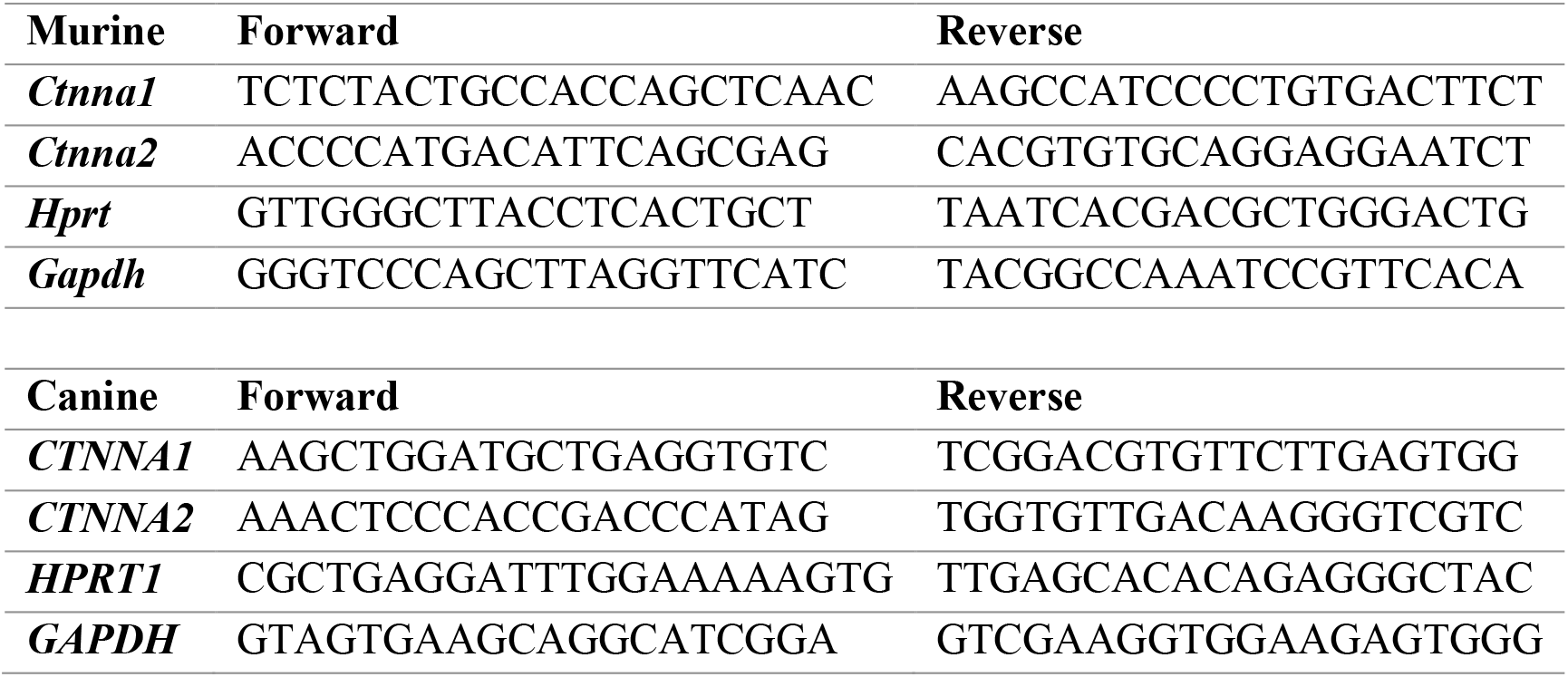

### Live cell imaging

Transfected α-catenin KD cells were trypsinized, centrifuged with growth medium, and then resuspended and pre-incubated in HBSS/Hepes at 37°C for 5mins. These cells were then plated on fibronectin coated coverslips held in a Chamlide CMS coverslip holder and live cell images were taken with a Zeiss LSM800 confocal microscope using a 63x 1.4NA objective at an interval of 20 seconds.

### Statistical analysis

All experiments were repeated at least twice on separate days using duplicates and triplicates on each day. All quantifications represent the mean ± standard error of the mean. Two-tailed unpaired Student’s *t* test or the Mann Whitney test was used for group comparisons whereas multiple group comparisons were performed by ANOVA followed by Tukey’s test as indicated in the figure legends. Differences were considered to be statistically significant from a p-value below 0.05.

## Acknowledgments

H.W. acknowledges support from the Israel Science Foundation (1738/17) and from the Rappaport Family Foundation. H.W. is an incumbent of the David and Inez Myers Career Advancement Chair in Life Sciences. Author contributions: A.M. performed most of the experiments and data analysis; E.N. performed cloning and assisted in generation of KD cell lines; S.M. the co-IP experiments; H.K. the western blots; M.A. the Real-time PCR and L.F. assisted with the immunostaining M.P.S and H.W. conceived the idea for the studies. H.W. supervised the studies. A.M. and H.W. wrote the manuscript. Competing interests: The authors have no competing interests.

