## Supplementary figures for "α-catenin links integrin adhesions to F-actin to regulate ECM mechanosensing and rigidity-dependence"

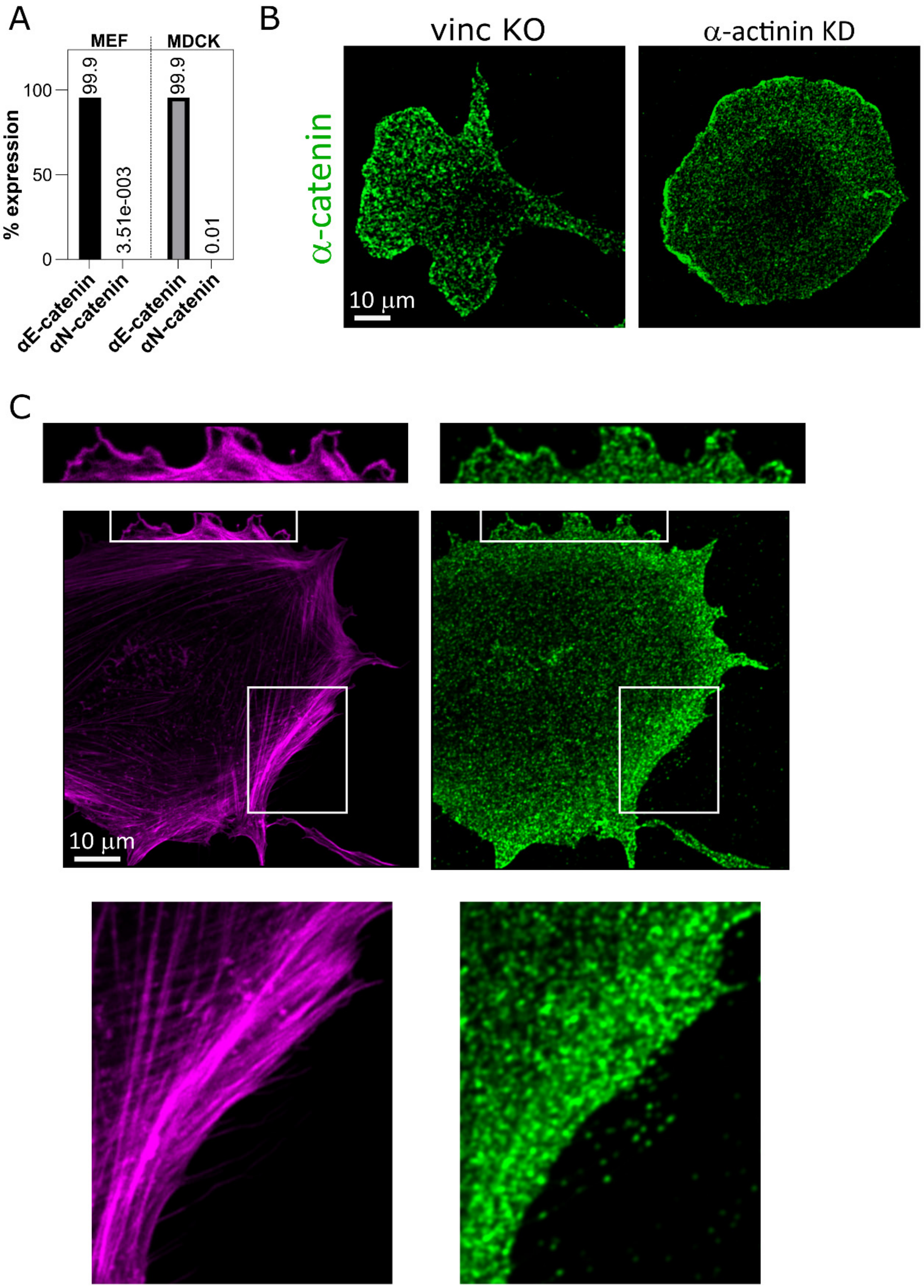


**Supplementary Fig. 1.** (A) RT-PCR analyses show the MEF and MDCK cells (used later in this study, see Fig. 5) express almost exclusively αE-catenin and not αN-catenin. (B) Immunostaining for α-catenin in vinculin-/- cells and α-actinin KD MEFs shows that its recruitment to the cell edge is not impaired by the absence of either of these proteins. (C) Immunostaining for α-catenin (green) in MEFs fixed 4 hours after plating on FN-coated glass shows co-localization with F-actin (magenta) on stress fibers (bottom zoom-in) as well as at the cell edge (top zoom-in). The brightness of the top rectangle in the zoomed-out image was enhanced for purpose of clarity.

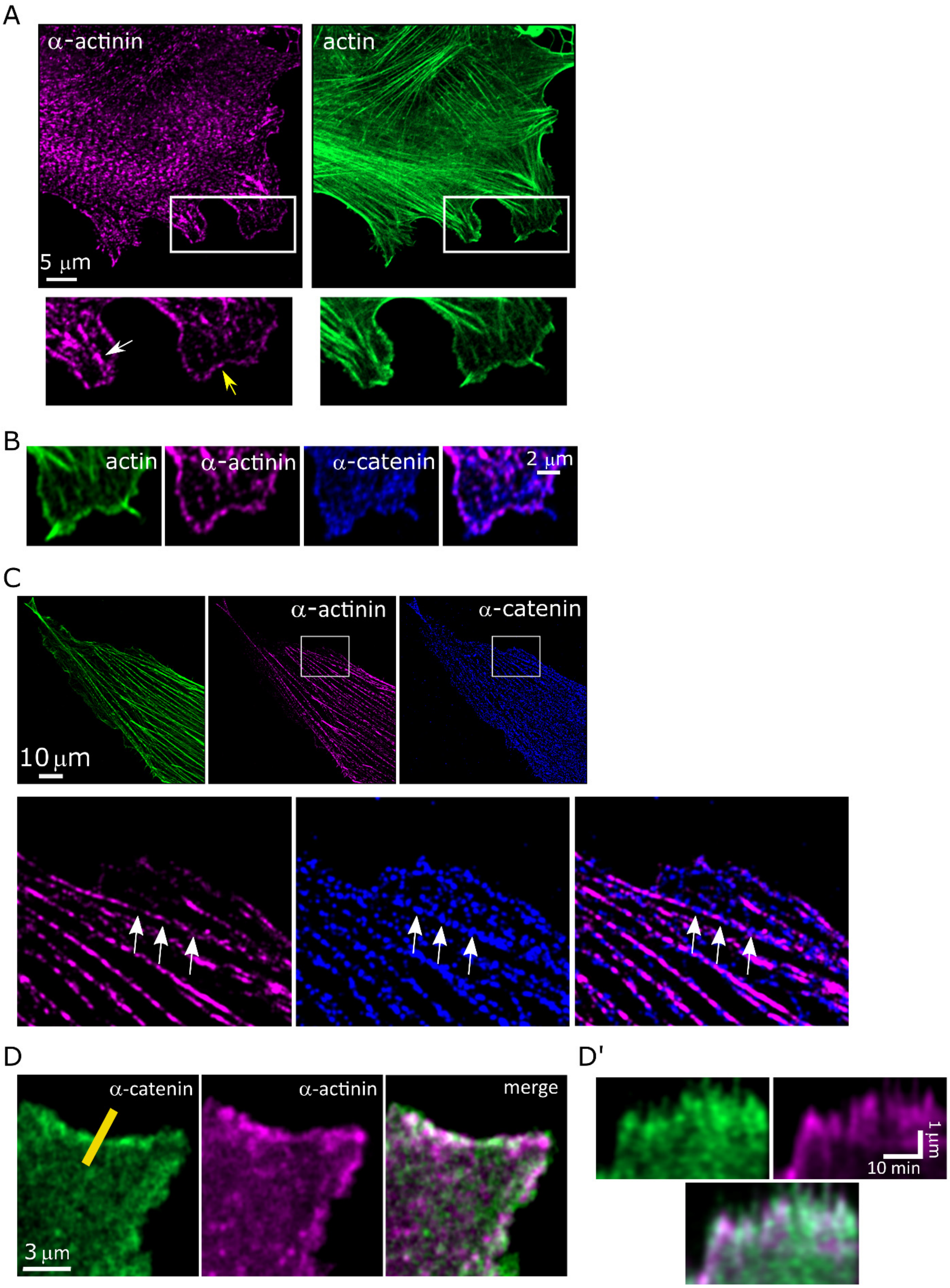


**Supplementary Fig. 2.** (A) Immunostaining in WT MEFs for α-actinin shows its co-localization after 15 min of spreading with F-actin at the cell edge (yellow arrow) as well as its localization in nascent adhesions (white arrow). (B) Zoom-in on the edge of a WT MEF cell co-stained for actin, α-actinin, α-catenin, showing lack of overlap of the latter two. (C) Immunostaining for α-actinin and α-catenin in WT MEFs shows their lack of overlap in actin stress fibers. Bottom row is the zoom-in of the boxes in the top row; arrows point to locations in which α-catenin is localized but α-actinin is not. (D) Frame from a movie of a cell expressing GFP-α-catenin and mCherry-α-actinin showing that two are not colocalized; (D') Kymographs taken from the yellow line shown in panel B.

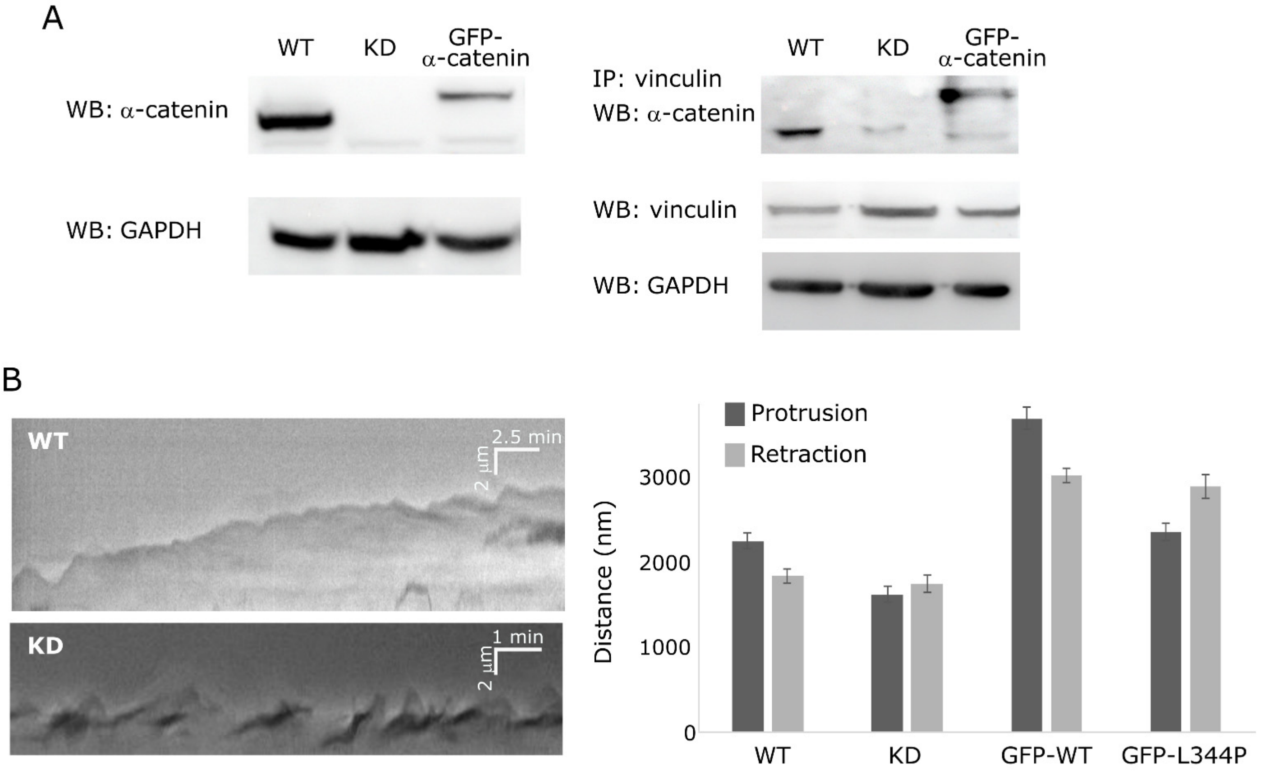


**Supplementary Fig. 3.** (A) Left: Immunoblot for α-catenin in lysates taken from WT MEFs, α-catenin KD MEFs, and α-catenin KD MEFs expressing WT GFP-α-catenin. Right: Immunoprecipitation for vinculin followed by immunoblot for α-catenin; below is a blot for vinculin from the cell lysates. (B) Left: Kymographs of the cell edge taken from time-lapse videos of early spreading by WT and α-catenin KD MEFs showing regular protrusion/retraction cycles in the former and extensive ruffling in the latter. Right: quantifications of the distances travelled during the protrusion and retraction phases by WT MEFs, α-catenin KD MEFs, α-catenin KD MEFs expressing WT GFP-α-catenin, and α-catenin KD MEFs expressing GFP-α-catenin L344P. WT and GFP-WT cells display longer protrusions than retractions, whereas KD and L344P cells do not, consistent with ruffling. N > 30 cycles from at least 8 cells in each case.

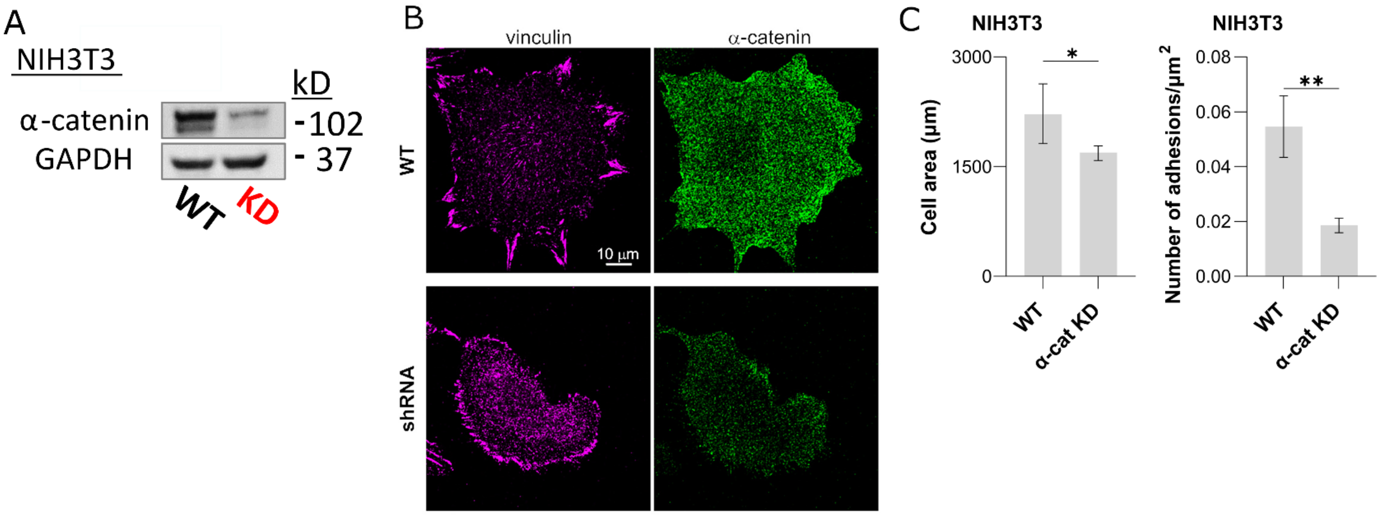


**Supplementary Fig. 4.** (A) Immunoblot for α-catenin showing KD in NIH3T3 cells. (B) Immunostaining for vinculin and α-catenin in WT and α-catenin KD NIH3T3 cells. (C) Quantification of cell area and number of mature adhesions per µm2 in WT and α-catenin KD NIH3T3 cells. N = 17 cells in each case. Statistical analysis for the NIH3T3 cell area and number of adhesions was performed with Student's t-test with Welch's correction (*, $p < .05$; **, $p < .01$; ***, $p < .001$, ****, $p < .0001$).

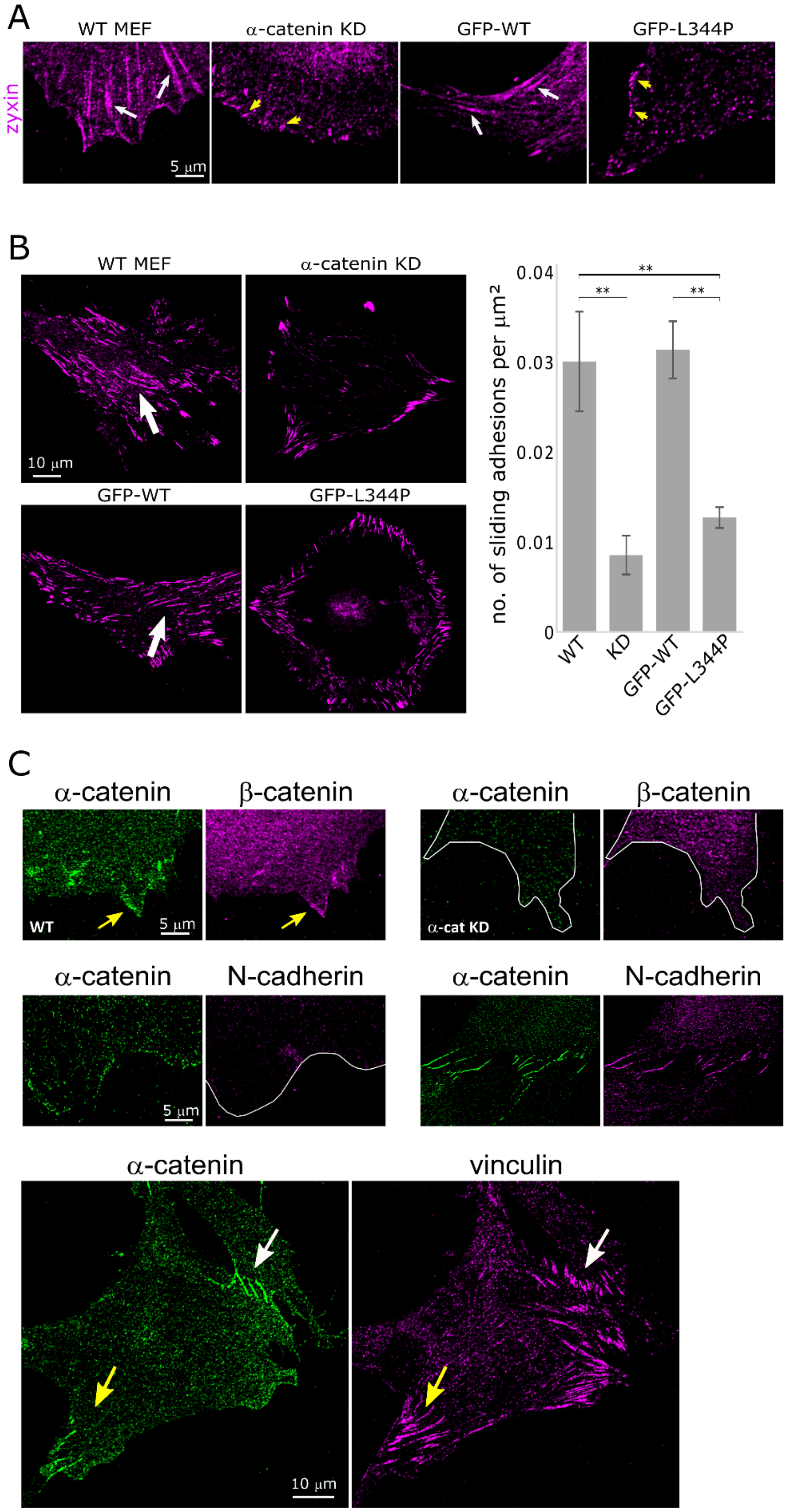


**Supplementary Fig. 5.** (A) Immunostaining for zyxin shows the presence of mature adhesions in WT MEFs and α-catenin KD cells expressing WT GFP-α-catenin (white arrows), but only presence of focal complexes in α-catenin KD cells and α-catenin KD cells expressing GFP-α-catenin L344P (yellow arrows). (B) Formation of sliding adhesions (arrows) on Matrigel-coated coverslips by WT cells, α-catenin KD cells, and α-catenin KD cells expressing WT GFP-α-catenin or GFP-α-catenin L344P (N>15 cells in each case). (C) Top: α-catenin and β-catenin localize at the cell edge, but β-catenin is missing from the edge upon α-catenin KD. Middle: N-cadherin is localized in cell-cell junctions (right) but not at the cell edge (left) in WT MEFs. Bottom: Mature FAs (yellow arrow) form in regions rich with α-catenin in cells that form α-catenin-rich cell-cell contacts (white arrow). Statistical analysis of the number of sliding adhesions was performed by ANOVA followed by Tukey's multiple comparisons test (*, $p < .05$; **, $p < .01$; ***, $p < .001$; ****, $p < .0001$).

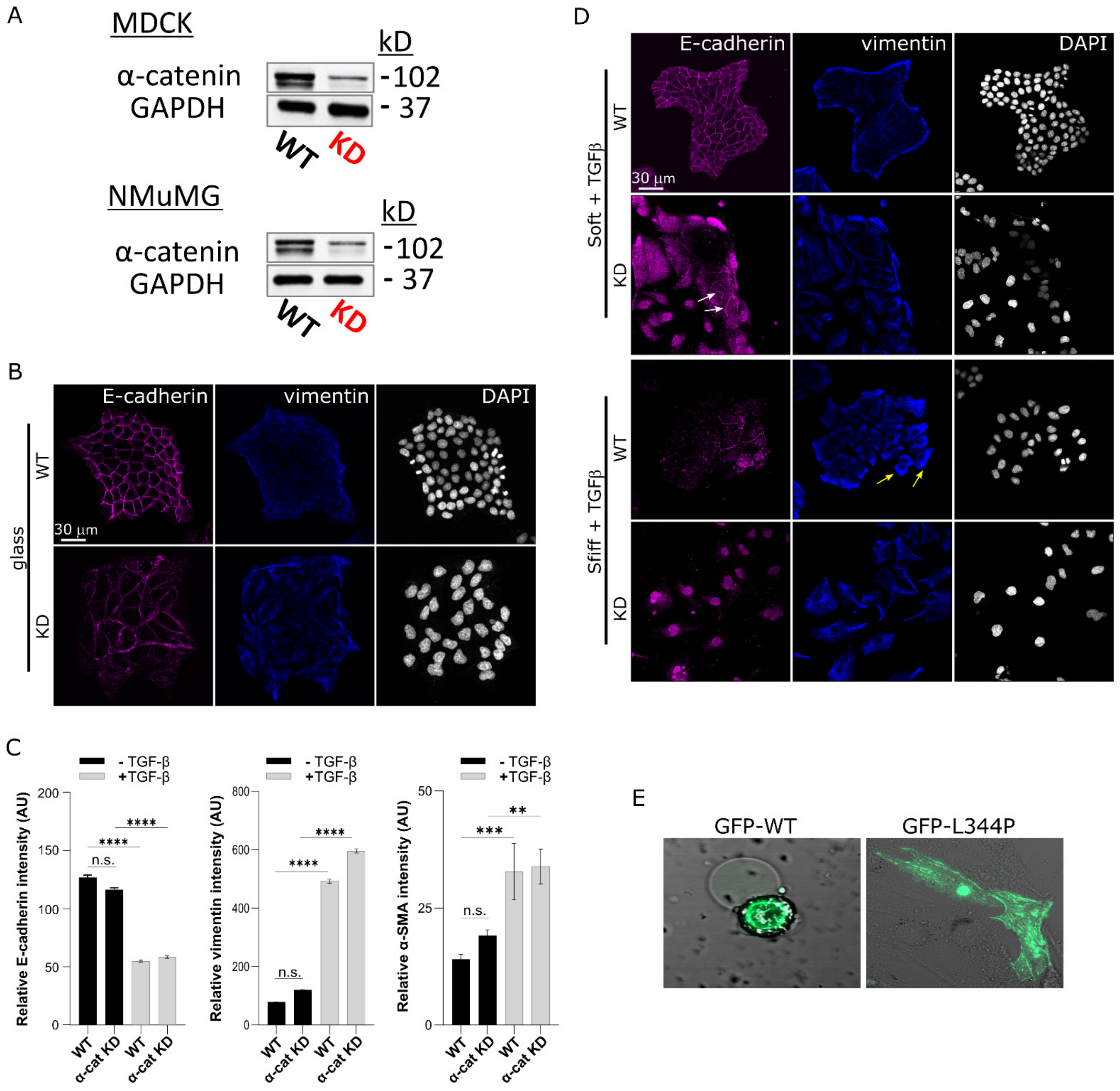


**Supplementary Fig. 6.** (A) Immunoblot for α-catenin showing KD in MDCK and NMuMG cells. (B) WT and α-catenin KD MDCK cells stained for E-cadherin and vimentin after 48h on FN-coated glass without stimulation for EMT. (C) Quantifications of E-cadherin, vimentin, and α-SMA intensity in WT and KD cells untreated or after 72h treatment with TGFβ. (D) WT and α-catenin KD MDCK cells stained for E-cadherin and vimentin after 72h incubation with TGFβ (10 ng/ml) on FN-coated soft (0.2kPa) and stiff (25kPa) matrices. N > 22 cells in each case. (E) α-catenin KD MEFs expressing WT GFP-α-catenin or GFP-α-catenin L344P plated for 24 hours on 0.2kPa FN-coated matrices. The L344P mutant localizes to actin bundles in central regions of the cells (similar to Fig. 3C,E). Statistical analysis comparing the relative intensities of the EMT markers was performed by ANOVA followed by Tukey's multiple comparisons test (*, $p < .05$; **, $p < .01$; ***, $p < .001$; ****, $p < .0001$).
